# Enhancing Optical Properties and Stability of DNA-Functionalized Carbon Nanotubes with Cryoprotectant-Mediated Lyophilization

**DOI:** 10.1101/2025.08.26.672410

**Authors:** Aceer Nadeem, Aidan Kindopp, Ella Junge, Maryam Rahmani, Daniel Roxbury

**Affiliations:** Department of Chemical Engineering, University of Rhode Island, Kingston, Rhode Island 02881, United States; School of Chemistry and Biochemistry, Georgia Institute of Technology, Atlanta, Georgia 30322, United States

**Keywords:** single walled carbon nanotubes, lyophilization, cryoprotectants, sensor stability, long-term storage, near infrared fluorescence

## Abstract

The long-term optical performance and stability of single-walled carbon nanotubes (SWCNTs) functionalized with single-stranded DNA are critical for their application in near-infrared (NIR) fluorescence biological sensing and imaging. However, the aggregation of such DNA-SWCNTs during storage presents a significant challenge. Here, we explored the use of lyophilization combined with various cryoprotectants to enhance the long-term stability and reconstitution of DNA-SWCNTs at room temperature. Five conventionally used cryoprotectants, including glucose, sucrose, mannitol, polyethylene glycol (PEG), and polyvinyl alcohol (PVA), were evaluated for their ability to maintain desired optical properties and prevent aggregation of SWCNTs through the process of lyophilization and reconstitution. Our results indicated that glucose and PEG, particularly in an 80:20 ratio by weight, provided the best performance, preserving NIR fluorescence and ensuring consistent reconstitution without significant aggregation. Further, *in vitro* studies using murine macrophages demonstrated that lyophilized SWCNTs with glucose-PEG protectants and then held at room temperature before subsequent reconstitution maintained stable intracellular optical performance, supporting their potential for long-term storage, ease of transport, and use in biomedical applications. These findings suggest that the optimized lyophilization protocol with specific cryoprotectant combinations can significantly improve the shelf life and reproducibility of SWCNT-based sensors, paving the way for their broader application in biological and clinical settings.

## Introduction

Single-walled carbon nanotubes (SWCNTs) are a class of nanomaterials characterized by their unique one-dimensional structure, described as a graphene sheet rolled seamlessly into a cylindrical shape, forming a hollow tube with an average diameter of about 1 nanometer. ^1^ This resulting structure exhibits remarkable mechanical, electrical, and optical properties.^2^ SWCNTs can be uniquely characterized by their chiral indices, which are defined by a specific pair of integers denoted as (n, m).^3^ With all of their atoms effectively on the surface of the nanomaterial, SWCNTs are highly sensitive to environmental variations^4^ and are ideal for a wide range of applications, particularly in the fields of biomedicine.^5^ SWCNTs exhibit intrinsic fluorescence within the near-infrared (NIR) spectrum,^6^ specifically in the wavelength range of 900 to 1400 nanometers. This spectral range is particularly significant because it falls within the tissue transparency window,^7^ a region where biological tissues exhibit minimal absorption and scattering of light. As a result, NIR fluorescence from SWCNTs can effectively penetrate biological tissues, making them highly suitable for applications in biological environments.^8^ Furthermore, the fluorescence emitted by SWCNTs in this range is photostable.^9^ These properties are crucial for long-term imaging^10^ and continuous monitoring, as they ensure consistent signal strength and reliability.^11^ Consequently, SWCNTs are ideal candidates for optical biosensors, particularly *in vivo* applications,^12^ where stable and durable sensing capabilities are essential for monitoring biological processes within living organisms.

Due to the non-polar nature of carbon atoms, SWCNTs are naturally hydrophobic. This leads to a strong tendency for SWCNTs to aggregate, making them difficult to work with in biological and aqueous environments.^8^ To address this challenge and enhance their usability, particularly in biomedical applications, SWCNTs can undergo non-covalent or covalent functionalization.^13,14,15, 16^ Noncovalent wrappings can include a variety of materials, ranging from surfactants^17^ to biocompatible molecules such as proteins^18^, and other biopolymers^19^ like PEG-lipid conjugates. By wrapping in these materials, the SWCNTs gain enhanced solubility and stability while maintaining their essential optical and electronic characteristics.^8^ For biological applications, SWCNTs are commonly non-covalently functionalized with single-stranded DNA (ssDNA), where the amphiphilic ssDNA wraps the SWCNT in a helical fashion, forming a DNA-SWCNT hybrid.^20, 21^ The unbounded hydrophobic nitrogenous bases of ssDNA adsorb the nucleic acid to the SWCNT surface while the hydrophilic backbone of the polymer stabilizes the DNA-SWCNT hybrid by facilitating interactions with polar water molecules.^22, 23, 24^ The wrapping and resultant stability of SWCNTs with DNA is dependent on the specific sequence of the DNA strand, in addition to pH, and other ions, small molecules, or macromolecules that may be in the immediate vicinity of the hybrid.^25, 26^ DNA-SWCNTs have been used in a wide variety of applications including for specific-biomarker recognition.^27, 28, 29, 30, 31^

The ability to store nanoparticles for an extended period while maintaining their stability has been an important topic in biological research for clinical applications. In the case of DNA-SWCNTs, the optimum storage and stability is crucial for their use in biomedical and sensing applications because any degradation or aggregation over time could impair their performance. Vacuum drying is the most basic form of nanoparticle storage, a process where the sample is heated to remove moisture. The nanoparticles go through one or more phase transformations during this process to end in a solid form,^32^ however, the method can cause significant aggregation.^33^ Supercritical fluids (SCF) that exceed the critical temperature and pressure, have also been used for nanoparticle storage because they possess both liquid and gas properties such as low viscosity, high density, and ability to act as a solute for insoluble solvents.^34, 35, 36^ However, this process involves thorough optimization of process parameters such as high temperature and pressure, has limited scalability, and further research is necessary to evaluate the biocompatibility of the method.^37^ The most commonly employed method for long-term nanoparticle storage is lyophilization, characterized by the freezing and subsequent removal of a solvent by sublimation under vacuum.^38^ Lyophilized nanoparticles maintain physical and chemical properties of the original product, have short reconstitution time, low residual moisture, long-term stability,^39, 40^ and scalable manufacturing.^41^ Muramatsu et al. demonstrated through an *in vivo* study that when lyophilized, mRNA lipid nanoparticles (mRNA-LNP) can be stored at room temperature for 12 weeks without the mRNA-LNP losing its high translatability.^42^ Lyophilized products are typically rehydrated before use, allowing them to regain their original properties with minimal loss of activity or efficacy.^43^

During the lyophilization process, chemical and physical stresses can be induced on the nanoparticles.^44^ To stabilize the nanoparticles, two types of protectants are utilized: cryoprotectants and lyoprotectants.^45^ The primary function of cryoprotectants is to protect the nanoparticles during the freezing process. Cryoprotectants cause water to melt at lower temperatures, helping to achieve regulation of the rates at which water transport, nucleation, and ice formation occur.^46^ Lyoprotectants stabilize nanoparticles during both the freezing and drying process.^45^ Typically, cryoprotectants can also act as lyoprotectants.^45^ Various sugars are commonly used as protectants used during the lyophilization such as mannitol, sucrose, and glucose due to their ability to be vitrified during freezing.^47^ Other, non-sugar protectants include dimethyl sulfoxide (DMSO), ethylene glycol, many polymers, and glycerol.^48^ The type and concentration of protectant, as well as the concentration of nanoparticles impacts the degree to which the nanoparticles are stabilized.^32^ In this study, we explored the use of various cryoprotectants for the long-term storage of DNA-SWCNTs. The primary objective was to enhance the stability of dispersed DNA-SWCNTs at room temperature, improve their re-dispersion after storage, and prevent unwanted aggregation, which is critical for maintaining their functionality. We systematically evaluated different cryoprotectants and optimized their ratios to identify the combination that provided the best performance in terms of intracellular stability and sensor efficacy over a four-week period. Our findings demonstrate that it is possible to store DNA-SWCNTs at room temperature for extended periods while preserving their intrinsic physical and optical properties. This advancement supports the long-term reliability of DNA-SWCNTs in various sensing applications, ensuring consistent performance over time.

## Results and Discussion

We synthesized DNA-SWCNTs by using standard probe-tip sonication and ultra-centrifugation, where single-stranded (GT)_30_ DNA was combined with HiPco SWCNTs, as detailed in the methods section. The (GT)_30_ DNA was selected as the test wrapping for this experiment due to its extensive use in the literature for non-covalent functionalization of SWCNTs,^49,50,51,52^ highlighting its relevance for various biosensing applications. Throughout the preparation process, meticulous attention was paid to ensure the production of a high-quality, monodispersed (GT)_30_-SWCNT dispersion, minimizing any potential for pre-existing aggregation.

Variations in SWCNT synthesis^53^ as well as DNA-SWCNT functionalization can often lead to inconsistencies in sample preparation from batch to batch. Moreover, long-term room temperature storage of these biopolymer dispersions can induce DNA degradation which leads to sample aggregation and a resulting decline of NIR fluorescence properties, which can pose significant challenges for reproducibility in resesrch.^54^ To address the inherent issues of long-term storage posed by the possibility of sample aggregation over time, lyophilization was tested as a potential method to mitigate modulations in fluorescence and achieve more consistent and reproducible results. We developed an optimized lyophilization protocol specifically designed to extend the long-term shelf life of SWCNTs and considerably reduce batch-to-batch variations. Our optimized protocol involved preparing a large, homogenous batch of DNA-SWCNT dispersion, which was then lyophilized. This approach ensured that each lyophilized batch originated from the same initial dispersion, thus reducing variability.

Following sample preparation, the DNA-SWCNT dispersion was flash frozen at -80°C for 60 minutes and then lyophilized for an additional 24 hours at 0.060 pa and -42°C to remove moisture and produce a powdered sample. The lyophilized DNA-SWCNT powder was subsequently reconstituted (“L+R”) in ultra-pure water by pipette mixing and compared to the pre-lyophilization (“Pre-L”) dispersion.

Figure 1a illustrates the NIR fluorescence comparison between the pre-lyophilized SWCNTs (Pre-L) and the lyophilized and then reconstituted SWCNTs (L+R). The spectrum was divided into four distinct bands to aid in data analysis and comparison. The NIR fluorescence spectrum shows a substantial decrease of approximately 45% in fluorescence intensity in the lyophilized sample (without the addition of CPs). The freeze-drying process is shown to generate a variety of stresses which can in turn impact nanoparticle stability.^55^ Figure 1b provides a schematic overview of how lyophilization of SWCNTs can lead to potential aggregation and thus loss of sample properties. The decrease in fluorescence intensity is also attributed to aggregation observed upon lyophilization of SWCNTs without any CPs as shown in SI Movie 1a and 1b. Figure 1c is a proposed schematic of the process used to optimize SWCNT reconstitution after lyophilization, highlighting how the use of cryoprotectants (CPs) can potentially mitigate aggregation and improve resuspension. For this purpose, five commonly used CPs from the pharmaceutical industry were evaluated for their lyophilization effectiveness with SWCNTs (Table S1, Figure S1)._56,57,58,59,60_

**Figure 1.**
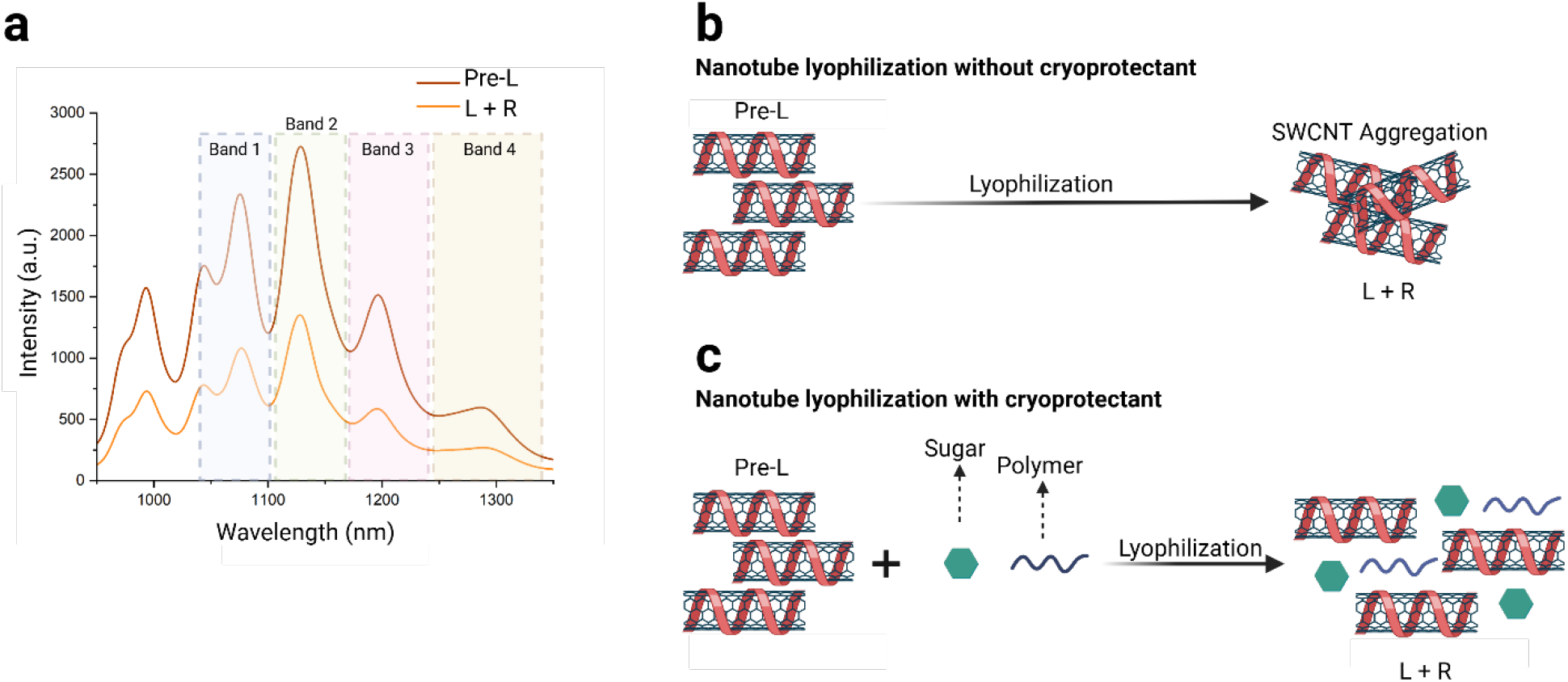
DNA-SWCNT lyophilization. (a) NIR fluorescence plot showing comparison between Pre-lyophilization (Pre-L) and lyophilization and reconstitution (L+R) of DNA-SWCNTs (no cryoprotectants added) taken under 730 nm excitation. (b) Lyophilization schematic indicating aggregation of SWCNTs (c) Proposed Lyophilization schematic detailing the process of lyophilization with added cryoprotectants (CPs).

Based on literature, three sugars, glucose, sucrose, and mannitol, as well as two polymers, polyethylene glycol (PEG) and polyvinyl alcohol (PVA) were chosen for investigation in this study. Only low molecular weights and weight percentages of polymers were examined as CPs as it was observed that higher molecular weights and weight percentages of polymers led to a significant aggregation of DNA-SWCNTs (data not shown). To achieve optimal performance, the CPs were incorporated with SWCNTs and systematically investigated for many factors, such as rapid reconstitution, optical response and long-term stability. Factors considered during final SWCNT cryoprotectant additive selection are mentioned in Table S2.

A freshly prepared DNA-SWCNT sample was added to 1 ml of each selected CPs to achieve a final SWCNT concentration of 5 mg-L^-1^ and the absorbance of each sample measured (Figure S2). No appreciable changes in the absorbance spectrum with respect to the as-dispersed SWCNT reference were observed for DNA-SWCNT samples upon addition to CPs, with the exception of PVA (Figure 2a, Figure S2e). A detailed table of absorbance change observed at 990nm in SWCNT (L+R) samples with respect to the as-dispersed SWCNT sample is shown in Table S3. Normalized absorbance data suggests glucose and sucrose show positive trends in absorbance recovery after reconstitution, indicating effective cryopreservation upon reconstitution (Figure S3). Mannitol and PEG also perform well upon reconstitution, whereas PVA shows a significant decrease in intensity with a degradation of nearly 57% in absorbance and sample quality upon reconstitution. The trends observed for changes in absorbance after reconstitution of lyophilized samples are the same for change in normalized NIR fluorescence of Band 2 for (L+R) samples with respect to the as-dispersed SWCNT sample (Figure 2b). Similar data for the other NIR fluorescence band intensities and the relative wavelength ranges considered during analysis of normalized data (Figure S4) can be found in Table S4.

**Figure 2.**
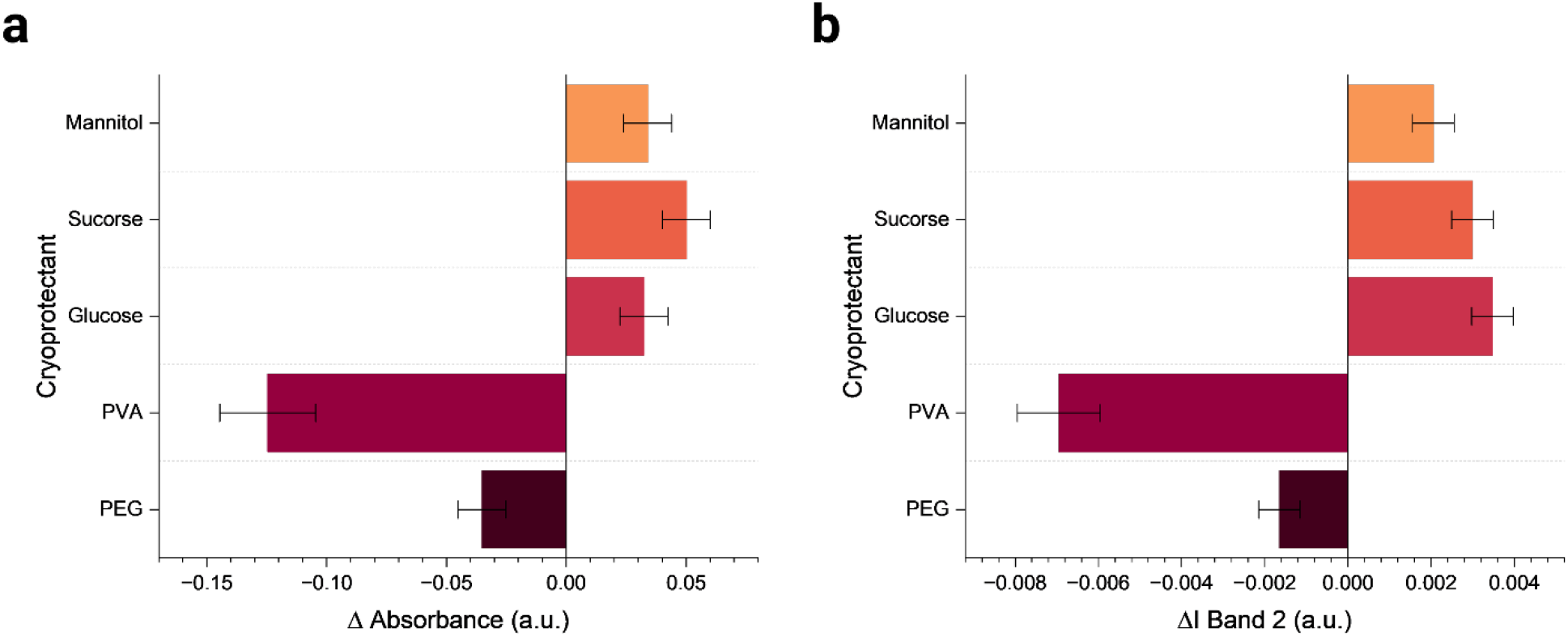
NIR characterization of lyophilization samples. (a) Change in absorbance of reconstituted lyophilized samples with CPs in comparison to as dispersed (GT)_30_ SWCNTs. (b) Change in band 2 peak intensity of reconstituted lyophilized samples with CPs in comparison to as dispersed (GT)_30_ SWCNTs. Error bars represent mean +/- st.dev. from n= 3 technical replicates.

A considerable decrease in fluorescence was noted with the PVA sample, exhibiting a significant 27% reduction in fluorescence compared to the as-dispersed sample. In contrast, the sample with PEG resulted in an 8% increase in fluorescence intensity. All sugar samples also performed well in NIR fluorescence quality checks with glucose giving the most consistent performance with a 13% increase in intensity. In summary, the data show considerable promise in performance and quality of samples for all CPs used except for PVA. The poor performance of PVA as a CP has also been reported for gold nanoparticles^61^ and, in our DNA-SWCNT system, can likely be attributed to strong interactions between PVA and the adsorbed (GT)_30_ ssDNA molecule which is preferentially complexing with the PVA polymer^62^, thus leading to SWCNT aggregation (Figure S5).

To reach an informed decision regarding the quality and performance of all DNA-SWCNT (L+R) samples, lyophilized powders of each sample were analyzed *via* Raman spectroscopy (Figure S6) and Fourier transform infrared (FTIR) spectroscopy (Figure S7) to elucidate quantitative structural information. Traditionally, Raman spectroscopy of SWCNTs focuses on analyzing the G-band (1585 cm^−1^), which scales linearly with the concentration of SWCNTs, and the D-band (1310-1350 cm^−1^), also known as the disorder or defect band, which quantifies defects on the nanotube surface.^63^ In this study, G-band sharpness obtained from Raman microscopy was utilized to quantify the crystalline structure^64^ of the DNA-SWCNT samples with cryoprotectant. The Raman spectrum for glucose- and sucrose-containing samples showed broad peaks, indicative of their amorphous nature. In contrast, the spectrum for PEG and PVA-containing samples exhibited well-defined sharp peaks, reflecting their strong crystalline structure.^65^ Notably, the PVA sample showed a significantly higher G/D ratio, which we speculate is due to the interaction between PVA and the DNA-SWCNTs, leading to aggregation^66^ and is further evidence for the formation of a PVA-DNA-SWCNT complex. Similar conclusions can be observed in the FTIR results, where the (GT)_30_ SWCNT control sample and PEG samples present well-defined peaks in contrast to the sugar samples that appear to have a slightly amorphous structure. A summary of the crystalline versus amorphous nature of each CP as well as more detailed FTIR spectral analysis can be found in Table S5. All CPs were further tested for in-vitro compatibility as shown in Figure S8 and S9. It was observed that PVA was not a good choice as CP for our system due to aggregation.

Based on all characterization results, glucose and PEG were selected as the most suitable CP candidates for the lyophilization of SWCNTs because glucose can form a stable amorphous matrix, and PEG provides additional colloidal stability without disrupting the glucose matrix nor inducing aggregation. In contrast, PVA displayed a high degree of aggregation when lyophilized with SWCNTs, negatively affecting reconstitution efficiency, as well as the absorbance and NIR fluorescence responses. Sucrose and mannitol were not pursued further due to complications regarding polymorphism which can induce batch to batch variability leading to stability issues, and other potential complications in downstream biomedical applications.^67,68^

Additional investigations into various weight ratios of glucose:PEG were conducted to identify the optimal combination for achieving the best long-term results as previous work has shown that combinations of multiple cryoprotectants have shown promise in exhibiting superior cryoprotection.^69^ Figure 3a and Figure 3b represent the pre-lyophilization absorbance spectroscopy and NIR fluorescence results for the different CP ratios of glucose and PEG tested. In the pre-lyophilization stage, no significant differences were observed across the CP weight ratios, except for the combination containing 0% glucose and 100% PEG. The different weight ratios were then lyophilized and reconstituted, and the absorbance spectra measured (Figure 3c). All samples containing CPs performed better than the reconstituted control (no CP) sample, except for the sample with 100% PEG. The NIR fluorescence spectra for all lyophilized and reconstituted samples were measured (Figure 3d), and a trend emerged that higher concentrations of PEG resulted in a decrease in the average fluorescence intensity. We hypothesize that this reduction in NIR fluorescence with increasing PEG concentration is due to a small yet non-negligible degree of aggregation of DNA-SWCNTs at higher PEG levels. The weight ratios of 80:20, 60:40, and 50:50 showed the best performance and were therefore selected for further long-term and *in vitro* cell studies. Although the sample with 100% glucose also exhibited satisfactory NIR fluorescence spectra, it was not chosen for any further investigations due to characterization results indicating the amorphous nature of the lyophilized DNA-SWCNT sample with glucose as cryoprotectant.

**Figure 3.**
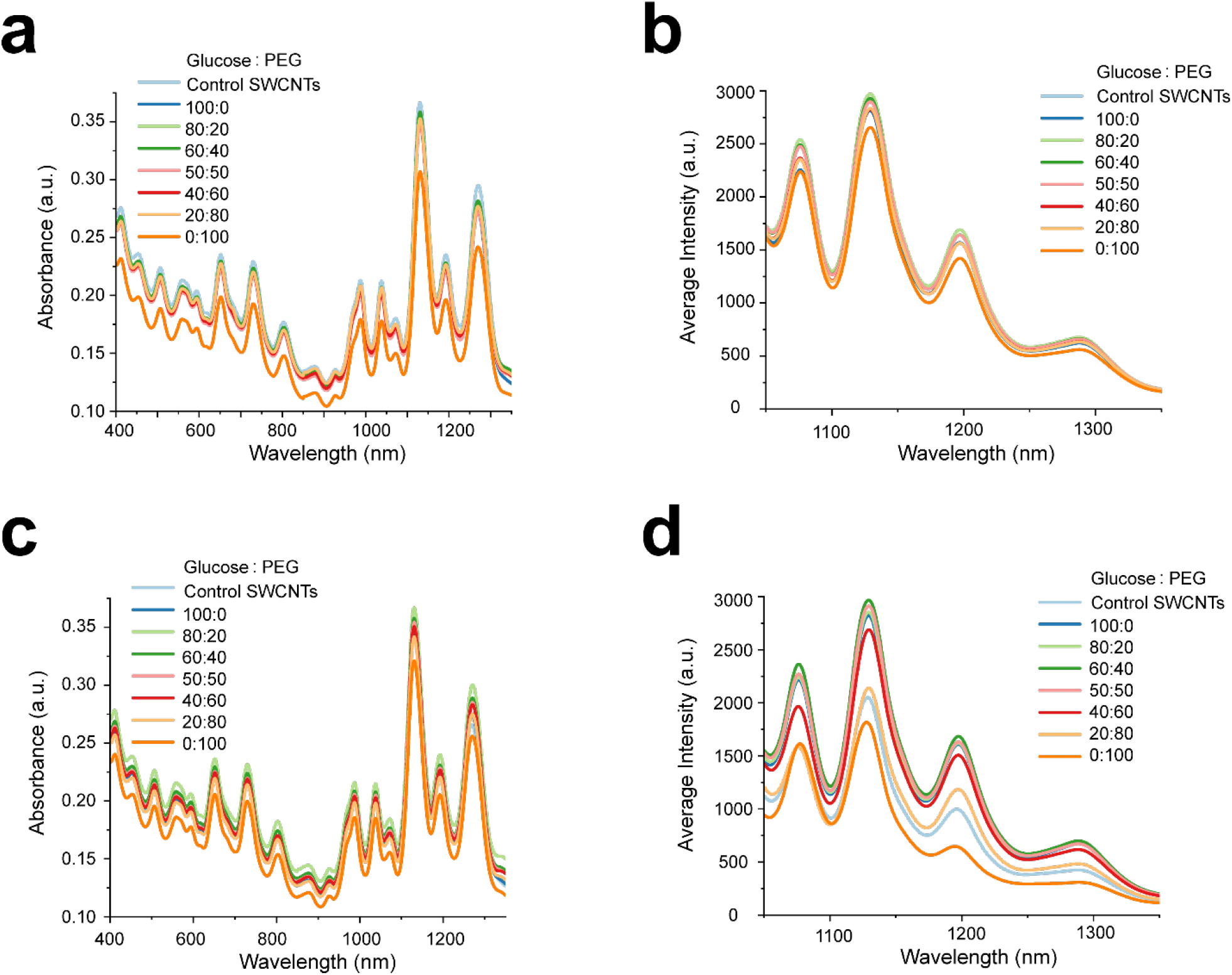
Evaluating differential weight ratios of glucose:PEG. (a) Absorbance spectroscopy plot comparing spectrum of DNA-SWCNT samples with different weight ratios of glucose:PEG pre-lyophilization and control SWCNTs with no CPs. (b) NIR fluorescence spectra of DNA-SWCNT samples with different ratios of glucose:PEG pre-lyophilization and control SWCNTs with no CPs (c) Absorbance spectroscopy spectra comparing DNA-SWCNT samples with different ratios of glucose:PEG post-lyophilization and reconstitution & control SWCNTs with no CPs. (d) NIR fluorescence plot of DNA-SWCNT samples with different ratios of glucose:PEG post-lyophilization & reconstitution and control SWCNTs with no CPs

We assessed the internalization and *in vitro* optical performance of the selected ratio L+R samples in live cultured cells. Macrophages (murine RAW 264.7), which are crucial components of the immune system and serve as the body’s first line of defense,^70^ were chosen and could be compared to our own and other groups’ previous studies.^28,61,68-71^ The cultured macrophages were incubated with 1 mg-L^-1^ of each selected ratio of glucose-PEG (L+R) and a GT_30_ SWCNT control sample (no CPs) for 30 minutes. Following this pulse incubation, the cells were thoroughly washed with 1x PBS (phosphate-buffered saline) and placed in fresh media for an additional 30 minutes before imaging at the 1-hour time point. Here, the as-dispersed DNA-SWCNTs were used as a standard control for all intracellular experiments. Figure 4a shows the intracellular broadband fluorescent images for all samples and distinct changes in both 1 hour and 24 hours can be observed. Figure 4b and c show NIR broadband fluorescence spectral response of the images in 4a. As expected, all glucose-PEG ratio samples performed better than the as-dispersed DNA-SWCNT control. The 80:20 and 60:40 glucose: PEG sample showed a larger increase in intensity relative to control, while the behavior of 50:50 sample was similar in nature to the as-dispersed SWCNTs. The increased fluorescence performance of these glucose and PEG ratios is attributed to increased cellular uptake. We speculate that including cryoprotectant molecules such as glucose and polyethylene glycol may increase nanoparticle uptake in phagocytic cells such as macrophages. Figure S10 shows increased fluorescence response of SWCNTs inside macrophages when incubated with glucose and PEG.

**Figure 4.**
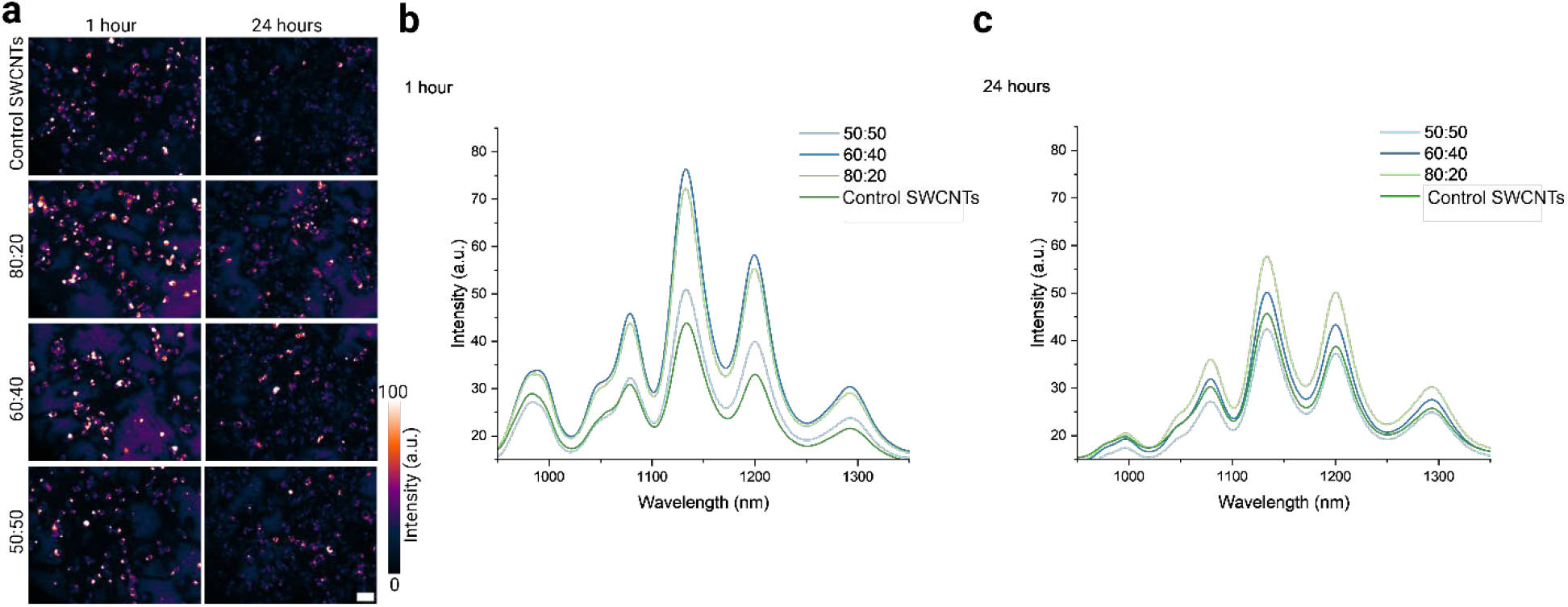
*In vitro* cell investigations of selected ratio L+R samples. (a) NIR broadband fluorescence (900-1400 nm) images of all selected ratio samples and the as dispersed control sample incubated with murine macrophages. Images have been globally contrasted to the brightest image. Scale bar =.15 µm (b) NIR fluorescence spectral response of cells incubated with selected SWCNTs (L+R) CP ratios and as-dispersed control sample at 1 hour and (c) 24 hours.

For long-term storage studies, the selected samples were analyzed for their NIR fluorescence response after storage for 2 or 4 weeks at three different temperatures: room temperature (RT, approximately 22°C), -20°C, and -80°C (Figure S11). For the samples stored at room temperature, both the 60:40 and 50:50 glucose-to-PEG ratios showed a notable decrease in NIR fluorescence, with more than 50% reduction and optical performance degradation in integrated NIR fluorescence intensity signal compared to the as-dispersed DNA-SWCNT (Figure 5a). In contrast, the 80:20 sample remained highly stable across all storage temperatures, including room temperature. We also observed that the NIR fluorescence at room temperature was quite stable between week 2 and week 4 for all the samples, meaning any degradation to quality of samples occurred within the first week. All samples stored at -20°C and -80°C exhibited similar responses, with no major differences observed (Figure S11) which is to be expected of lyophilized components under cold-chain storage. Interestingly, the 80:20 glucose-to-PEG DNA-SWCNT sample remained highly stable for a period of 12-months at room temperature as compared to the as-dispersed DNA-SWCNT sample, showing limited signs of degradation (figure 5b). The 60:20 and 50:50 ratios could not be tested as they did not properly resuspend after 12 months storage at room temperature, likely due to general sample degradation and aggregation.

**Figure 5.**
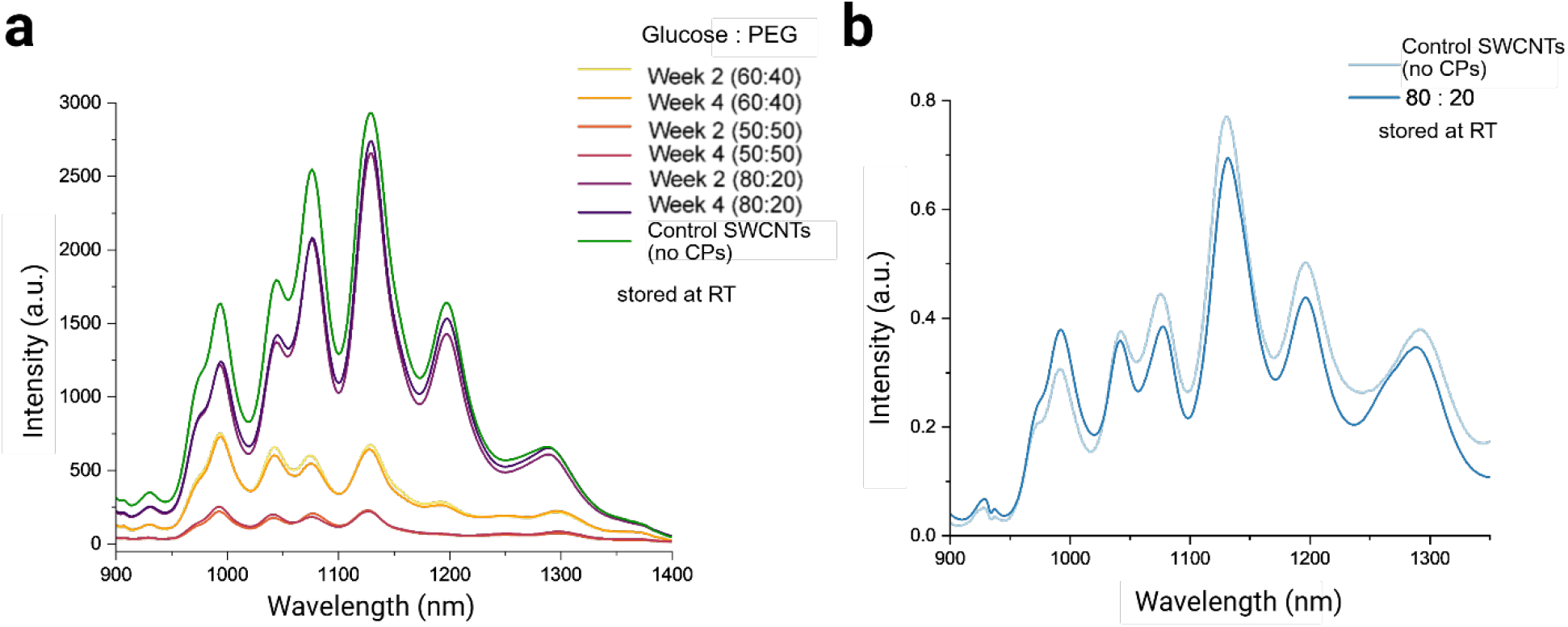
Long-term stability assessment of lyophilized DNA-SWCNTs stored at room temperature. (a) NIR fluorescence stability of lyophilized and reconstituted samples compared to control (as-dispersed) SWCNTs stored at room temperature for 2 and 4 weeks. (b) NIR fluorescence stability (normalized) of lyophilized and reconstituted (L+R) samples compared to control (as dispersed) SWCNTs stored at room temperature for 12 months.

Finally, to assess the ability of lyophilized and reconstituted DNA-SWCNTs to retain their sensing abilities, their response to hydrogen peroxide was examined and compared to as-dispersed DNA-SWCNT sample (Figure S12). Notably, the DNA-SWCNT L+R sample with 80:20 glucose:PEG ratio still responded to the addition of 2 mM hydrogen peroxide in a similar fashion as that of the as-dispersed DNA-SWCNT sample.

## Conclusion

The objective of this study was to systematically evaluate different CPs as potential additives to DNA-SWCNT samples to achieve more consistent, reproducible, and optimized outcomes with long-term stability. Five CPs were investigated, along with various ratios of the most effective candidates. Glucose and PEG were selected as the primary CPs due to their optimized optical performance, stability, and compatibility in *in vitro* cell studies. Further research showed that the 80:20 weight ratio of glucose:PEG offered the most optimized response for both intracellular optical performance and applications targeting the sensing of specific biomolecules. The 80:20 glucose:PEG ratio proved to be the most stable, performing best overall, and could be stored at room temperature without degrading the DNA wrapping or stability for up to 12 months. Lyophilizing SWCNTs specifically with 80:20 glucose:PEG ratio not only enhances practical applicability for long term storage but potentially revolutionizes storage and long-term handling of biopolymer suspended SWCNTs specifically for biomedical applications like biosensing and diagnostic tool development. This could in turn lead to more reliable and efficient technologies, significantly impacting healthcare outcomes by enabling more precise detection and monitoring.

## Acknowledgements

This work was supported by the National Science Foundation (CAREER Award #1844536 and #2231621) and the University of Rhode Island College of Engineering. The confocal Raman data were acquired at the RI Consortium for Nanoscience and Nanotechnology, a URI College of Engineering core facility partially funded by the National Science Foundation EPSCoR, Cooperative Agreement #OIA-1655221. Research was made possible by the use of equipment available through the Rhode Island Institutional Development Award (IDeA) Network of Biomedical Research Excellence from the National Institute of General Medical Sciences of the National Institutes of Health under grant #P20GM103430 through the Centralized Research Core facility. Schematics were created using BioRender.com software.

## TOC Graphic

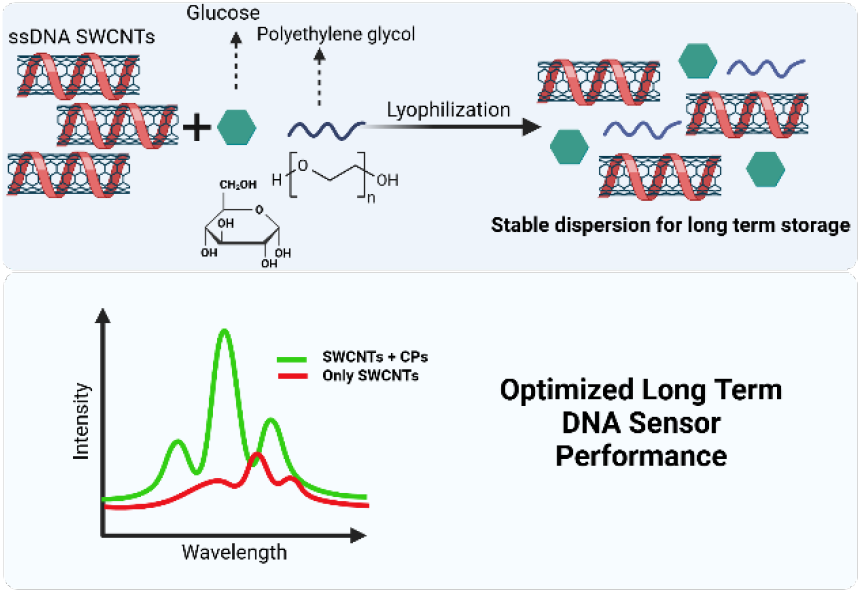

