## Supporting Information for "Enhancing Optical Properties and Stability of DNA-Functionalized Carbon Nanotubes with Cryoprotectant-Mediated Lyophilization"

by

*Aceer Nadeem[1,2], Aidan Kindopp[1], Ella Junge[1], Maryam Rahmani[1], and Daniel Roxbury[1]**

[1]Department of Chemical Engineering, University of Rhode Island,

Kingston, Rhode Island 02881, United States

[2]School of Chemistry and Biochemistry, Georgia Institute of Technology,

Atlanta, Georgia 30322, United States

### Methods and Materials

**DNA-SWCNT Sample Preparation.** To create monodispersed ssDNA wrapped SWCNTs (DNA–SWCNTs), 1 mg of HiPCO-SWCNTs was added to 2 mg of $(GT)_{30}$ oligonucleotide (Integrated DNA Technologies) in 1 mL of 0.1 M NaCl (Sigma-Aldrich). Each sample was ultrasonicated using a 1/8″ tapered microtip for 30 min at 40% amplitude in an ice bath (Sonics Vibracell VCX-130; Sonics and Materials). The resulting suspensions were ultra-centrifuged (Beckman Optima MAX-XP) for 30 min at 250,000 *g* and 4°C, and the top ∼80% of the supernatant was collected.

**Near-Infrared Fluorescence Microscopy.** A hyperspectral NIR fluorescence microscope, similar to a previously detailed system,[76] was used to obtain all hyperspectral fluorescence data. A 730 nm excitation laser source was reflected onto the sample stage of an Olympus IX-73 inverted microscope equipped with a LCPlan N, 20x/0.45 IR objective by Olympus, U.S.A. The resulting fluorescence emission was passed through a volume Bragg grating and collecting with a 2D InGaAs array detector by Photon Etc. (Montreal, Canada) to generate spectral image stacks. Live cell samples were mounted on a stage top incubator by Okolab, to maintain 37°C and 5% $CO_2$ cell culture conditions throughout the imaging procedure. All hyperspectral cubes, fluorescence images and transmitted light images were background corrected and processed in MATLAB.

**Confocal Raman Microscopy.** All Raman spectroscopy data were acquired using an inverted WiTec Alpha300R confocal Raman microscope (WiTec, Germany) equipped with a Zeiss Epiplan-NEOFLUAR 100x/1.3 Oil Pol, Oil immersion, objective, a 785nm laser (20 mW output measured at the sample), and a UHTS 300 spectrograph (300 lines/mm grating) coupled with an Andor DR32400 CCD detector (-61C, 1650 x 200 pixels). Multiple point spectrums were scanned, and spectra were obtained using a 15s integration time and 100 accumulations per spectrum to construct hyperspectral images. Background subtraction and cosmic ray removal were performed using a polynomial function in WiTec Project 5.2 software. Hyperspectral data were extracted and processed using custom codes written with MATLAB.

**Cell Culture.** RAW 264.7 TIB-71 cell line from ATCC (Manassas, VA, USA) was cultured under standard incubation conditions at 37°C and 5% CO2. “D-10” cell culture media containing sterile filtered high-glucose DMEM with 10% heat inactivated FBS, 2.5% HEPES, 1% L-glutamine, 1% penicillin/streptomycin, and 0.2% amphotericin B (all from Gibco) was used for cell culture. Cells were passaged every 2-3 days and used for experiments.

**Sample Preparation for Optical Microscopy.** For all 20x *in vitro* NIR fluorescence imaging experiments, the cells were plated, in triplicate, at an initial concentration 5.26 x $10^4$ cells/$cm^2$ on 35 mm glass-bottom microwell dishes (MatTek) and allowed to culture overnight. To dose the cells with DNA-SWCNT samples, the culture media was removed and replaced with 2 mL of 1 mg-$L^{-1}$ $GT_{30}$-SWCNTs diluted in D10 cell culture media and incubated for 30 minutes to allow cell internalization. The SWCNT-containing media was then removed, the cells were washed twice with sterile PBS (Gibco), followed by the addition of fresh media. All time points were defined with respect to this step.

**Cryoprotectant Solution Preparation (Sugars).** 10 wt. % solutions of glucose, sucrose, and mannitol were made by dissolving 10 g of each sugar in 100 mL of ultra-pure water. The solution was set to spin at 160 rpm for 1 hour until all sugar was dissolved. The solutions were then passed through a 0.22 μm filter, covered with parafilm and kept in the refrigerator for experiments. Sugar solutions were only used for up to 4 days for experiments to minimize bacterial contamination.

**Cryoprotectant Solution Preparation (Polymers).** 0.25 wt. % solutions of polyethylene glycol (MW ~ 1500) and polyvinyl alcohol (MW ~ 10,000) were made in ultra-pure water. The solution was set to spin at 160 rpm for 2 hours until all polymer was dissolved, covered with parafilm and kept at room temperature for experiments.

**Lyophilization of SWCNT.** SWCNT samples were added to 1 mL of each cryoprotectant solution to make up a final SWCNT concentration of 5 mg-L. The solutions were then flash frozen in a -80°C cryo-freezer (ThermoFisher) for 60 minutes. The frozen samples were then added to a lyophilizer at 0.060 Pa and -42°C for 24 hours. Lyophilized powders were then stored at respective storage temperatures.

**Statistical Analysis.** OriginPro 2022b was used to perform all statistical analyses. All data either met assumptions of statistical tests performed (i.e., equal variances, normality, etc.) or were transformed to meet assumptions before any statistical analysis was caried out. Statistical significance was analyzed using Two-Sample t-test or one way ANOVA where appropriate. Testing of multiple hypotheses was accounted for by performing one-way ANOVA with Tukey's posthoc test.

**Table S1.** List of cryoprotectants investigated in the study with their respective molecular weights and weight percentages used in all experiments

| Cryoprotectant | Molecular Weight / Weight Percent |
|---|---|
| Polyethylene glycol | MW – 1500 / 0.25 wt.% |
| Polyvinyl alcohol | MW – 10,000 / 0.25 wt.% |
| Glucose | 10 wt.% |
| Sucrose | 10 wt.% |
| Mannitol | 10 wt.% |

**Table S2.** Evaluated parameters during the selection process of cryopreservation.

| Parameter |
|---|
| Re-dispersibility |
| Long term stability – time* |
| Long term stability – storage temp* |
| Degree of aggregation |
| Degree of crystallinity |
| Change in NIR-fluorescence signal |
| *In-vitro* signal stability |

**Table S3.** Detailed table describing absorbance change observed at 990nm in SWCNT (L+R) samples with respect to the as dispersed SWCNT sample.

| CP Sample | Absorbance change (a.u.) | Trend vs. Control |
|---|---|---|
| PEG | -0.0351 | Decrease |
| PVA | -0.1246 | Large Decrease |
| Glucose | 0.0324 | Increase |
| Sucrose | 0.0501 | Increase |
| Mannitol | 0.034 | Increase |

**Table S4.** Detailed near infrared peak analysis for all fluorescence bands present in the SWCNT samples analyzed for lyophilization experiments.

| Band # | λ Range | $GT_{30}$ SWCNTs | Glucose | Mannitol | PEG | PVA | Sucrose |
|---|---|---|---|---|---|---|---|
| Band 1 | 1020-1080 | 0.021616 | 0.023403 | 0.024534 | 0.017199 | 0.015474 | 0.021972 |
| Band 2 | 1081-1140 | 0.025117 | 0.028586 | 0.027167 | 0.023463 | 0.018168 | 0.028111 |
| Band 3 | 1141-1250 | 0.018599 | 0.018599 | 0.018261 | 0.018391 | 0.016146 | 0.018137 |
| Band 4 | 1251-1350 | 0.00768 | 0.008978 | 0.007675 | 0.00893 | 0.008115 | 0.009192 |

**Table S5.** Detailed FTIR analysis on crystalline and amorphous structures of SWCNTs lyophilized with different cryoprotectants investigated.

| Cryoprotectant | St. Deviation. | Intensity Range | Structure Type |
|---|---|---|---|
| PVA | 8.24 | 60.38 | Crystalline |
| Glucose | 1.60 | 17.40 | Amorphous |
| Mannitol | 4.74 | 29.77 | Mixed |
| PEG | 7.20 | 57.40 | Crystaline |
| Sucrose | 5.83 | 29.11 | Mixed |

**Calculation Method for St. Deviation**

For a given sample (e.g., PVA), you have a series of intensity values across all wavenumbers. The standard deviation (SD) is calculated as:

$$SD = \sqrt{\frac{1}{N-1}\sum_{i=1}^{N}(x_i - \bar{x})^2}$$

where:

$x_i$ = individual intensity value at the i-th wavenumber

$\bar{x}$ = mean intensity value across all wavenumbers for that sample

$N$ = total number of data points (wavenumbers)

The standard deviation represents variability in peak intensities across three replicate FTIR measurements for each cryoprotectant used. A higher standard deviation indicated greater fluctuation between samples, suggesting less uniform molecular interactions or structural consistency in the sample. Lower values reflect more reproducible vibrational features and thus a more stable molecular environment. The intensity range describes the overall amplitude of the FTIR absorbance within the region of interest in the wavenumber. It describes how strongly the sample absorbs infrared light in that particular region, which directly corresponds to differences in bond vibrations, crystallinity as well as structural order. Larger intensity ranges will generally be associated with a higher degree of crystallinity while smaller values indicate amorphous structure.

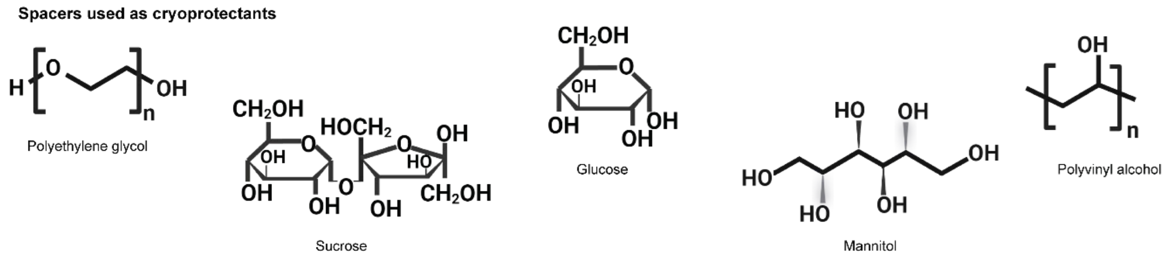


**Figure S1. Cryoprotectants for lyophilization.** Schematics showing chemical structures of cryoprotectants (CPs) that were investigated in this study.

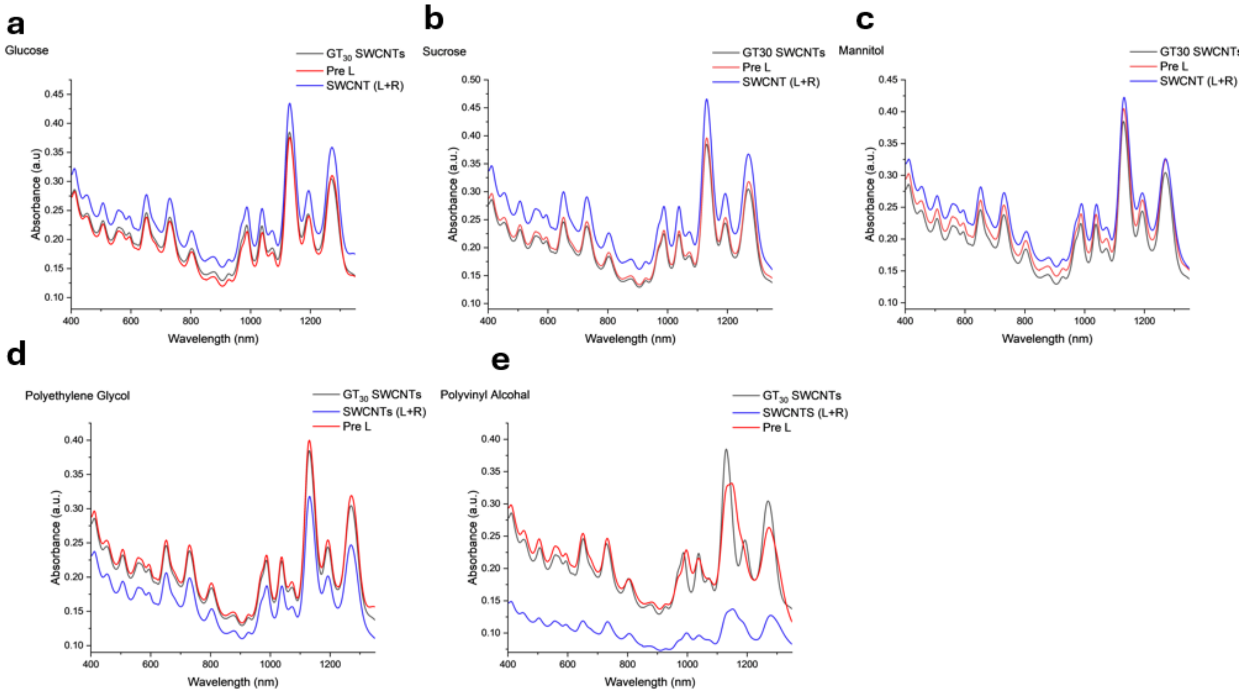


**Figure S2. Absorbance characterization of lyophilized DNA-SWCNTs.** Absorbance spectroscopy plots comparing spectrum of SWCNT samples with all CPs investigated pre-lyophilization (Pre L) and reconstituted lyophilized SWCNTs (L+R) for (a) glucose, (b) sucrose, (c) mannitol, (d) polyethylene glycol, and (e) polyvinyl alcohol.

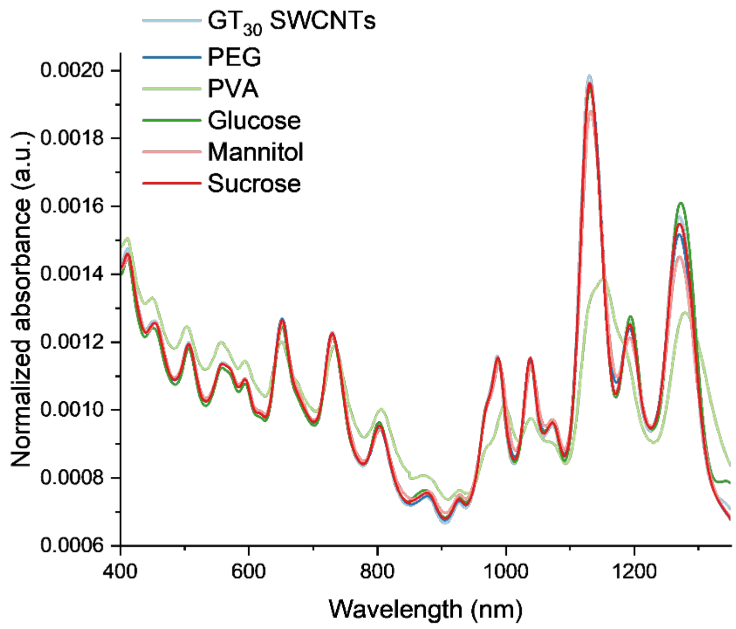


**Figure S3.** Normalized absorbance of all SWCNT (L+R) samples and the as-dispersed $GT_{30}$ SWCNT sample.

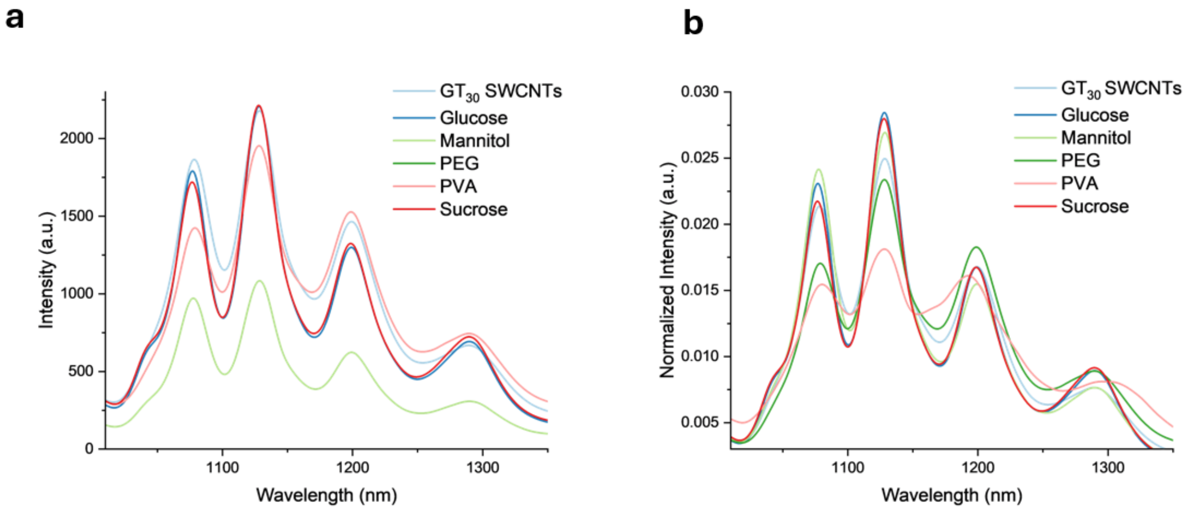


**Figure S4.** NIR fluorescence plot of DNA-SWCNT samples with all CPs investigated upon reconstitution after lyophilization and compared to as dispersed SWCNTs as (a) absolute intensity and (B) normalized intensity.

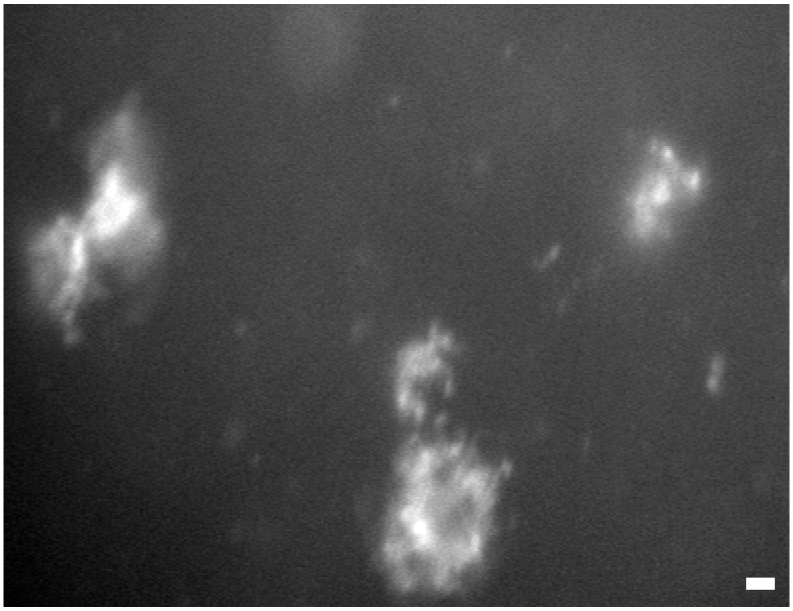

**Figure S5**. NIR broadband fluorescence image (900-1400 nm) showing substantial aggregation in DNA-SWCNT L+R PVA sample (Size of scale bar is 15µm.)

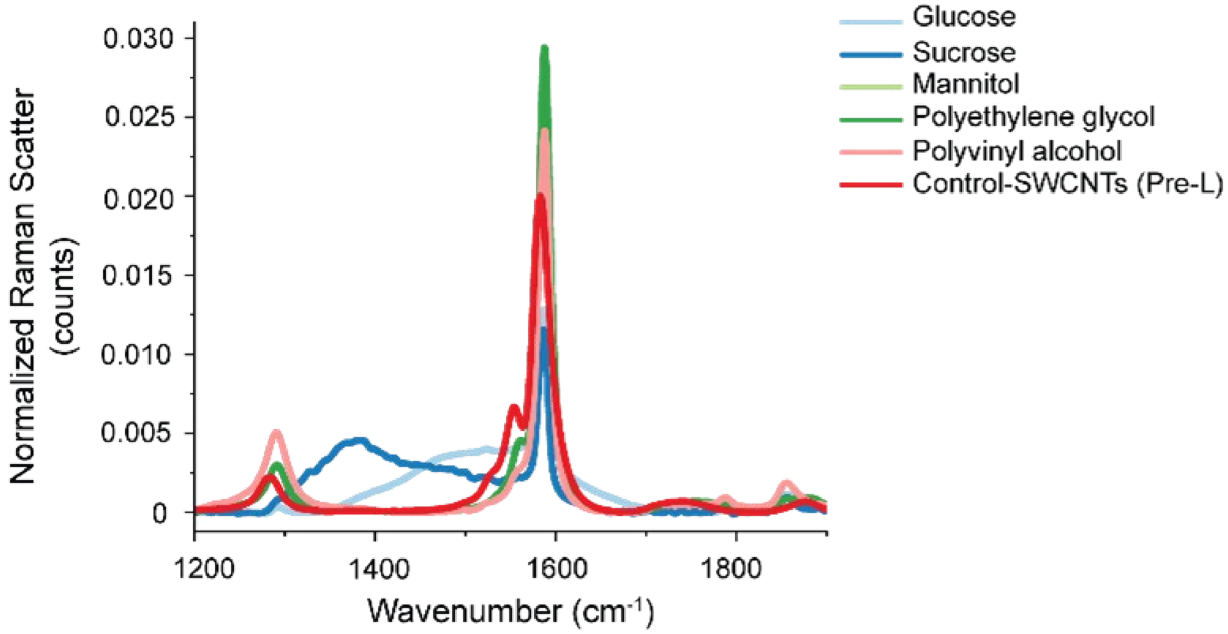


**Figure S6. Raman characterization of lyophilized DNA-SWCNTs.** Raman scatter spectra showing comparison between lyophilized samples and the control DNA-SWCNT sample (as produced, non-lyophilized sample).

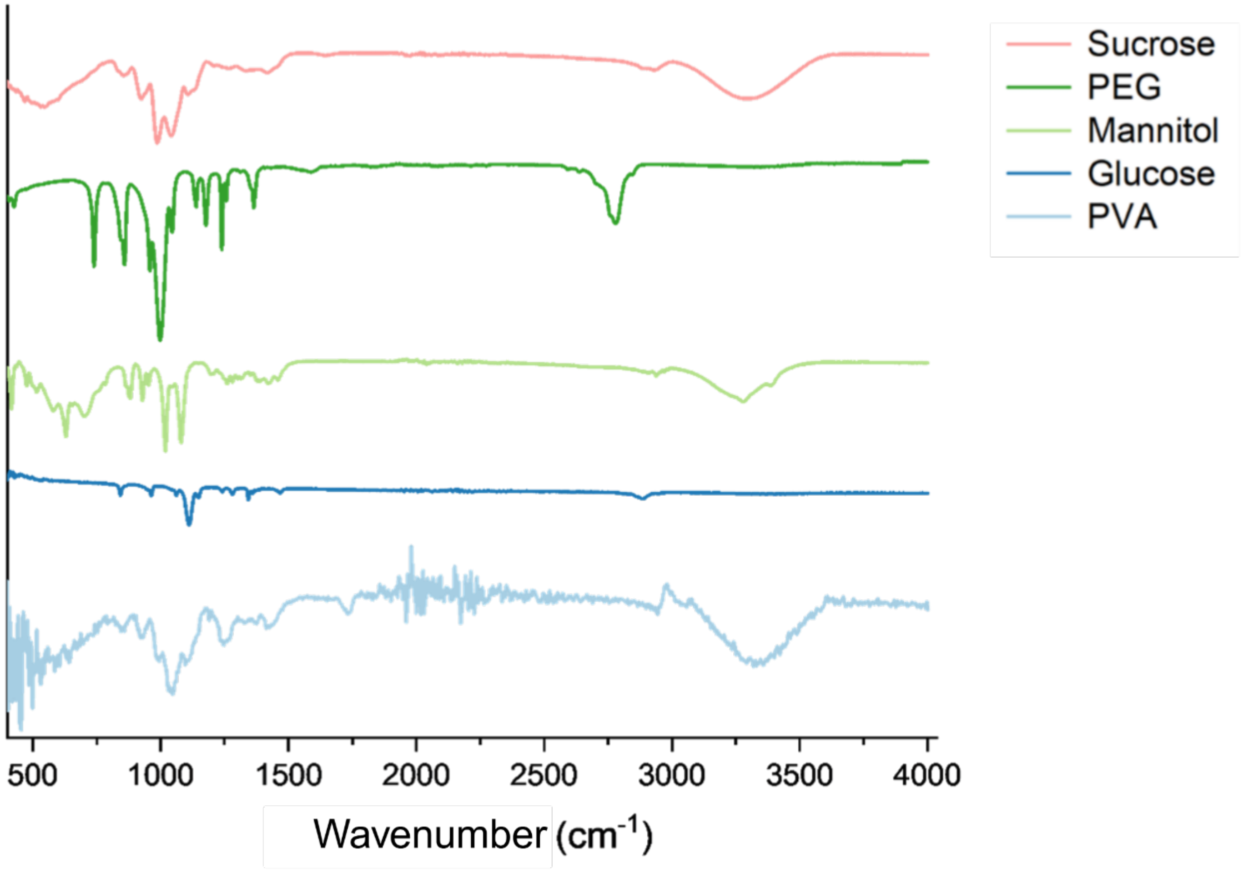


**Figure S7.** Fourier Transform Infrared Spectroscopy (FTIR) comparison of all lyophilized DNA-SWCNTs powders.

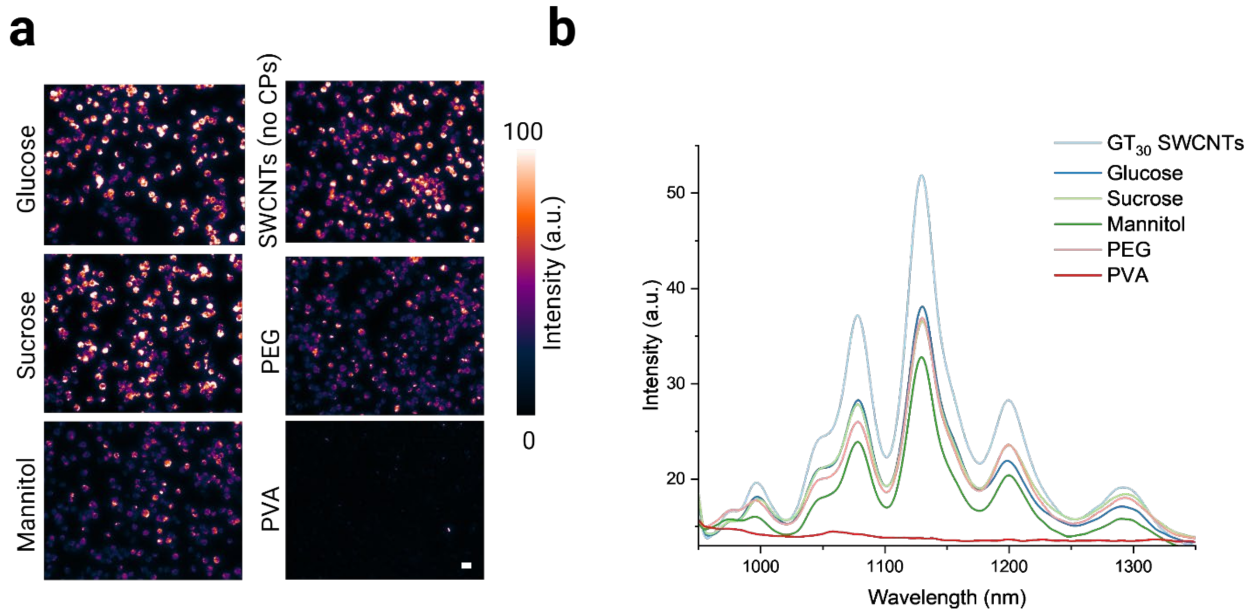


**Figure S8.** ***In vitro*** **cell investigations of DNA-SWCNT L+R samples.** (a) NIR broadband fluorescence (900-1400 nm) images of all DNA-SWCNT L+R samples incubated with murine macrophages. Images have been globally contrasted to the brightest image. Scale bar =.15 µm (b) Intracellular NIR fluorescence spectrum for all conditions as shown in (a).

Figure S8 shows NIR broadband fluorescence images (i.e. integrated fluorescence intensity from 900-1400 nm) and spectral response from all DNA-SWCNT L+R samples acquired at the 1-hour timepoint. As expected, glucose, sucrose, and PEG samples performed similar in response but displayed lower NIR fluorescence intensity than the as-dispersed DNA-SWCNT control. The mannitol sample experienced a decrease in intensity as compared to the other CPs. Finally, the PVA sample displayed limited internalization into cells, with most DNA-SWCNTs becoming aggregated and either getting removed in the washing step or getting stuck to the cell membrane as depicted in Figure S11b. Spectral analyses of such samples were disregarded as we chose to focus only on DNA-SWCNTs that internalized into the cells and gave the best results.

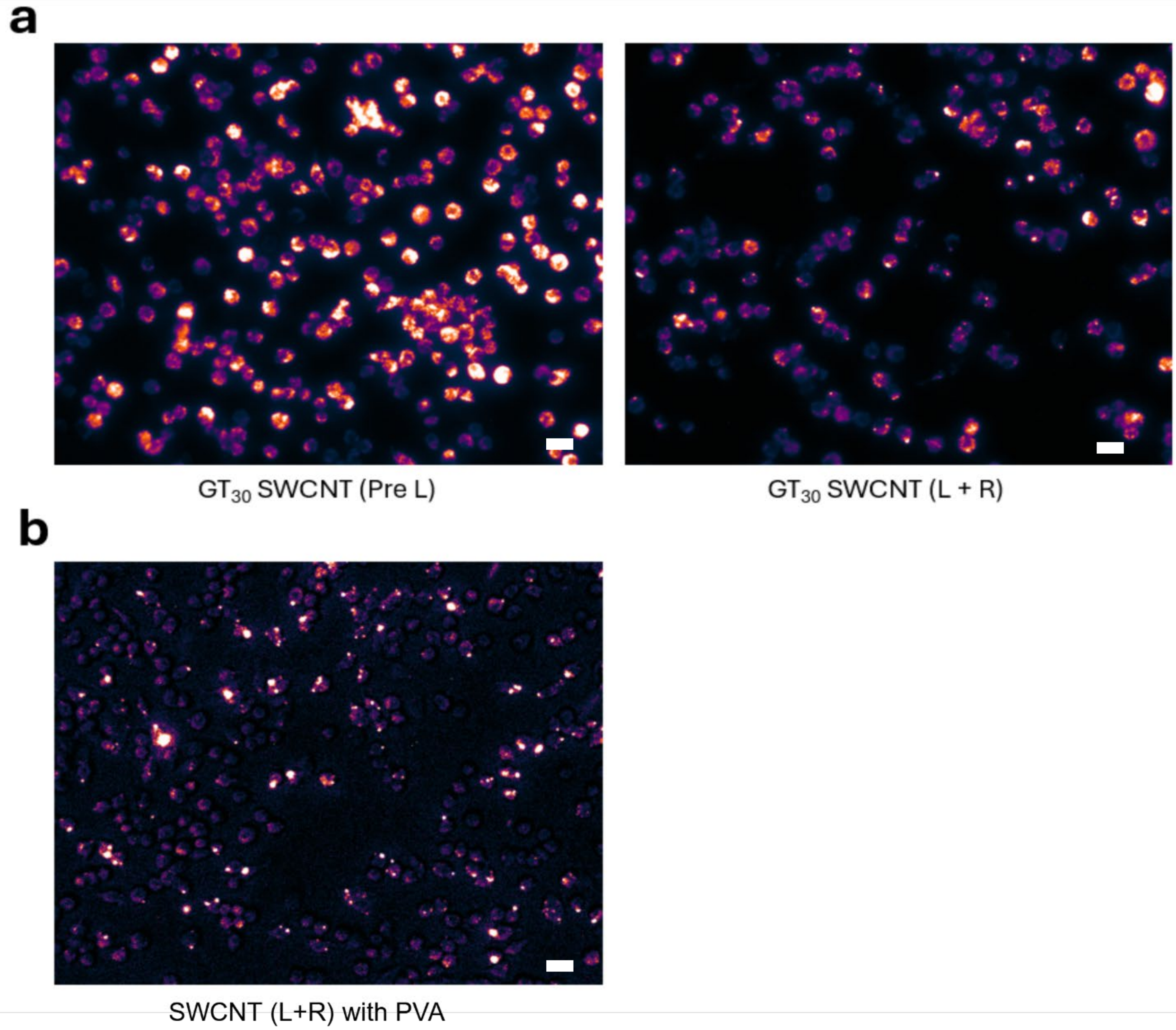


**Figure S9.** Intracellular NIR fluorescence broadband images of (a) $GT_{30}$ SWCNT sample before (as dispersed) and after L+R without the addition of any CPs. (b) Visible aggregation of DNA-SWCNT L+R sample with PVA. Image was contrasted to show aggregation and cells. (Size of scale bar is 15µm.)

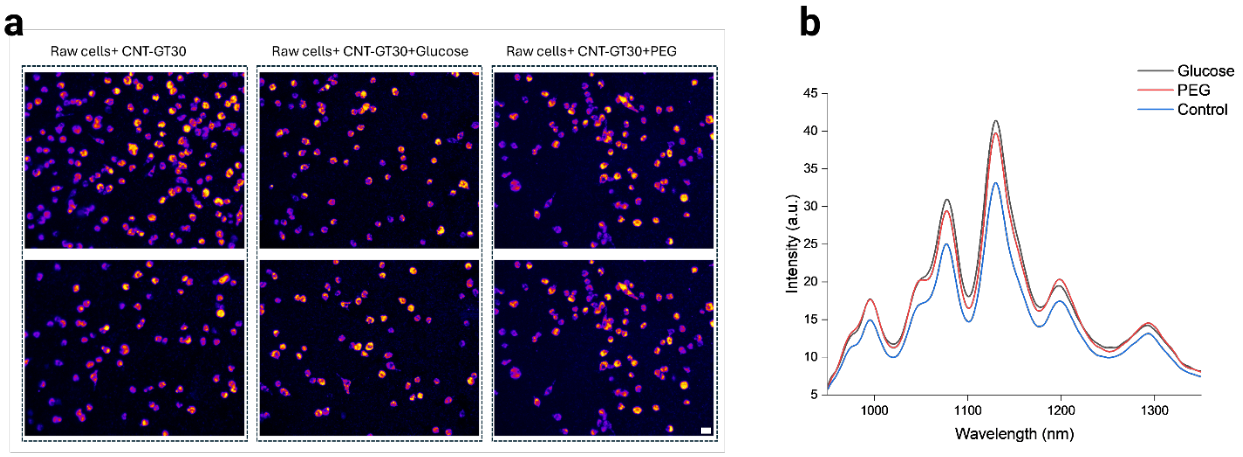


**Figure S10. Further *In vitro* cell investigations to observe enhanced optical response of glucose and PEG additives to SWCNTs.** (a) NIR broadband fluorescence (900-1400 nm) images of all DNA-SWCNT incubated with murine macrophages in presence of PEG and glucose. Images have been globally contrasted to the brightest image. Scale bar =.15 µm (b) Intracellular NIR fluorescence spectrum for all conditions as shown in (a).

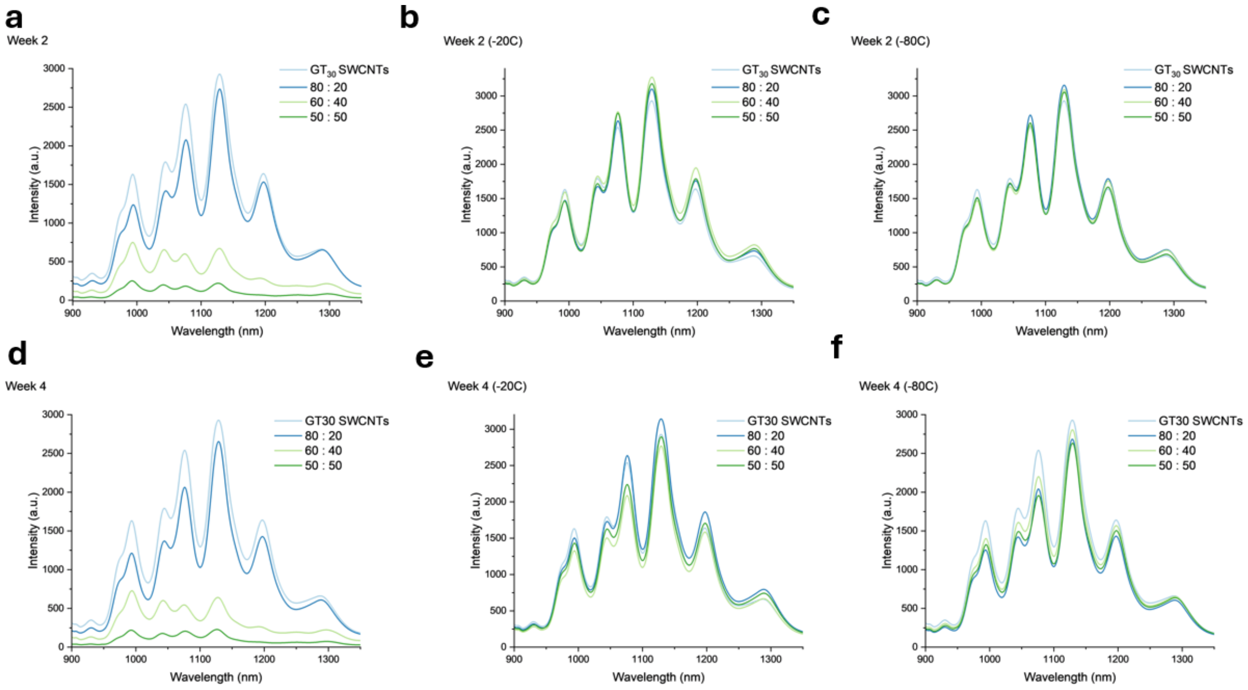


**Figure S11. Long term stability investigation.** Investigations into the long-term stability of chosen glucose:PEG ratios for 2 weeks at (a) room temperature, (b) -20°C, or (c) -80°C and for 4 weeks at (c) room temperature, (d) -20°C, or (e) -80°C.

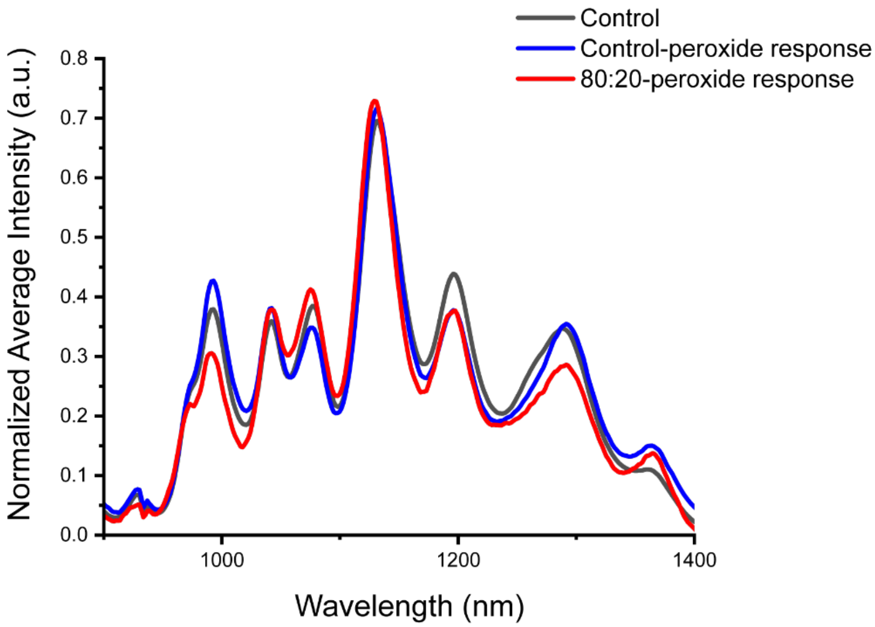


**Figure S12.** Near infrared fluorescence response comparison between control DNA-SWCNT (as dispersed) sample and the 80:20 glucose:PEG ratio L+R sample upon the addition of 2 mM hydrogen peroxide.
